# IFNβ and TNFα cooperate to induce a STAT1-independent antiviral and immunoregulatory program via non-canonical STAT2 and IRF9 pathways

**DOI:** 10.1101/273623

**Authors:** Mélissa K. Mariani, Pouria Dasmeh, Audray Fortin, Elise Caron, Mario Kalamujic, Alexander N. Harrison, Dacquin M. Kasumba, Sandra Cervantes-Ortiz, Espérance Mukawera, Adrian W.R. Serohijos, Nathalie Grandvaux

## Abstract

IFNβ typically induces an antiviral and immunoregulatory transcriptional program through the activation of ISGF3 (STAT1, STAT2 and IRF9) transcriptional complexes. The response to IFNβ is context-dependent and is prone to crosstalk with other cytokines, such as TNFα IFNβ and TNFα synergize to drive a specific delayed transcriptional program. Previous observation led to the hypothesis that an alternative STAT1-independent pathway involving STAT2 and IRF9 might be involved in gene induction by the combination of IFNβ and TNFα. Using genome wide transcriptional profiling by RNASeq, we found that the costimulation with IFNβ and TNFα induces a broad antiviral and immunoregulatory transcriptional program independently of STAT1. Additionally, STAT2 and IRF9 are involved in the regulation of only a subset of these STAT1-independent genes. Consistent with the growing literature, STAT2 and IRF9 act in concert to regulate a subgroup of these genes. Unexpectedly, STAT2 and IRF9 were also engaged in specific independent pathways to regulate distinct sets of IFNβ and TNFα-induced genes. Altogether these observations highlight the existence of distinct previously unrecognized non-canonical STAT1-independent, but STAT2 and/or IRF9-dependent pathways in the establishment of a delayed antiviral and immunoregulatory transcriptional program in conditions where elevated levels of both IFNβ and TNFα are present.

## INTRODUCTION

Interferon (IFN) β plays a critical role in the first line of defense against pathogens, particularly viruses, through its ability to induce a broad antiviral transcriptional response in virtually all cell types ^1^. IFNβ also possesses key immunoregulatory functions that determine the outcome of the adaptive immune response against pathogens ^1, 2^. IFNβ acts through binding to the IFNAR receptor (IFNAR1 and IFNAR2) leading to Janus kinases (JAK), JAK1 and Tyk2, mediated phosphorylation of signal transducer and activator of transcription (STAT) 1 and STAT2, and to a lesser extent other STAT members in a cell-specific manner ^3, 4^. Phosphorylated STAT1 and STAT2 together with IFN Regulatory Factor (IRF) 9 form the IFN-stimulated gene factor 3 (ISGF3) complex that binds to the consensus IFN-stimulated response element (ISRE) sequences in the promoter of hundreds of IFN stimulated genes (ISGs) ^5^. Formation of the ISGF3 complex is considered a hallmark of the engagement of the type I IFN response, and consequently the requirement of STAT1 in a specific setting has become a marker of the engagement of type I IFN signaling ^3, 6^. However, in recent years this paradigm has started to be challenged with accumulating evidence demonstrating the existence of non-canonical JAK-STAT signaling mediating type I IFN responses ^4, 7^.

Over the years, *in vitro* and *in vivo* studies aimed at characterizing the mechanisms and the functional outcomes of IFNβ signaling were mostly performed in relation to single cytokine stimulation. However, this unlikely reflects physiological settings, as a plethora of cytokines is secreted in a specific situation. As a consequence, a cell rather simultaneously responds to a cocktail of cytokines to foster the appropriate transcriptional program. Response to IFNβ is no exception and is very context-dependent, particularly regarding the potential cross-talk with other cytokines. IFNβ and TNFα exhibit context-dependent cross-regulation, but elevated levels of both cytokines are found during the host response to pathogens, including virus and bacteria, and also in autoinflammatory and autoimmune diseases ^8^. While the cross-regulation of IFNβ and TNFα is well studied, the functional cross-talk between these two cytokines remains poorly known and is limited to the description of a synergistic interaction ^9-12^. Indeed, costimulation with IFNβ and TNFα was found to drive a specific delayed transcriptional program composed of genes that are either not responsive to IFNβ or TNFα separately or are only responsive to either one of the cytokine ^10, 11^.

The signaling mechanisms engaged downstream of the costimulation with IFNβ and TNFα remained elusive, but it is implicitly assumed that the fate of the gene expression response requires that both IFNβ-and TNFα-induced signaling pathways exhibit significant cross-talk. Analysis of the enrichment of specific transcription factors binding sites in the promoters of a panel of genes synergistically induced by IFNβ and TNFα failed to give a clue about the specificity of the transcriptional regulation of these genes ^10^. Recently, analysis of the induction of *DUOX2 and DUOXA2* genes, which belong to the category of delayed genes that are remarkably induced to high levels in response to the combination of IFNβ and TNFα, led to the hypothesis that STAT2 and IRF9 activities might segregate in an alternative STAT1-independent pathway that could be involved in gene induction downstream of IFNβ and TNFα ^12^. Further validation was awaited to confirm the existence of this STAT1-independent response and the extent to which it is involved in the regulation of a specific delayed transcriptional program induced by the combination of IFNβ and TNFα.

In the present study, we aimed to characterize the transcriptional profile of the delayed response to IFNβ and TNFα in the absence of STAT1 and evaluate the role of STAT2 and IRF9 in the regulation of this (Inline) advantage of STAT1-deficient cells, we found that the synergistic action of IFNβ and TNFα induces a broad delayed antiviral and immunoregulatory transcriptional program independently of STAT1. We also report that STAT2 and IRF9 are differentially involved in the regulation of distinct subsets of genes induced by IFNβ and TNFα While IFNβ and TNFα act in part through the concerted action of STAT2 and IRF9, specific sets of genes were only regulated either by STAT2 or IRF9. These findings highlight the existence of distinct previously unrecognized non-canonical STAT2 and/or IRF9-dependent pathways that mediate the synergistic action of IFNβ and TNFα.

## RESULTS

### Distinct induction profiles of antiviral and immunoregulatory genes in response to IFNβ, TNFα and IFNβ+TNFα

First, we sought to determine the induction profile of a selected panel of immunoregulatory and antiviral genes in response to IFNβ+TNFα in comparison to IFNβ or TNFα alone. A549 cells were stimulated either with IFNβ TNFα or IFNβ+TNFα for various times between 3-24h and the relative mRNA expression levels were quantified by qRT-PCR. Analysis of the expression of the selected genes revealed distinct profiles of response to IFNβ TNFα or IFNβ+TNFα (**Figure 1**). *IDO*, *DUOX2*, *CXCL10*, *APOBEC3G, ISG20 and IL33* exhibited synergistic induction in response to IFNβ+TNFα compared to IFNβ or TNFα alone. Expression in response to IFNβ+TNFα increased over time, with maximum expression levels observed between 16 and 24h. While *NOD2* and IRF1 induction following stimulation with IFNβ+TNFα was also significantly higher than upon IFNβ or TNFα single cytokine stimulation, they exhibited a steady-state induction profile starting as early as 3h. *MX1* and *PKR*, two typical IFNβ-inducible ISGs, were found induced by IFNβ+TNFα similarly to IFNβ alone and were not responsive to TNFα. *CCL20* responded to IFNβ+TNFα with a kinetic and amplitude similar to TNFα, but was not responsive to IFNβ alone. *IL8* expression was not induced by IFNβ but was increased by TNFα starting at 3h and remained steady until 24h. In contrast to other genes, *IL8* induction in response to IFNβ+TNFα was significantly decreased compared to TNFα alone. Overall, these results confirm that induction of a subset of antiviral and immunoregulatory genes is greatly increased in response to IFNβ+TNFα compared to either IFNβ or TNFα alone.

**Figure. 1:**
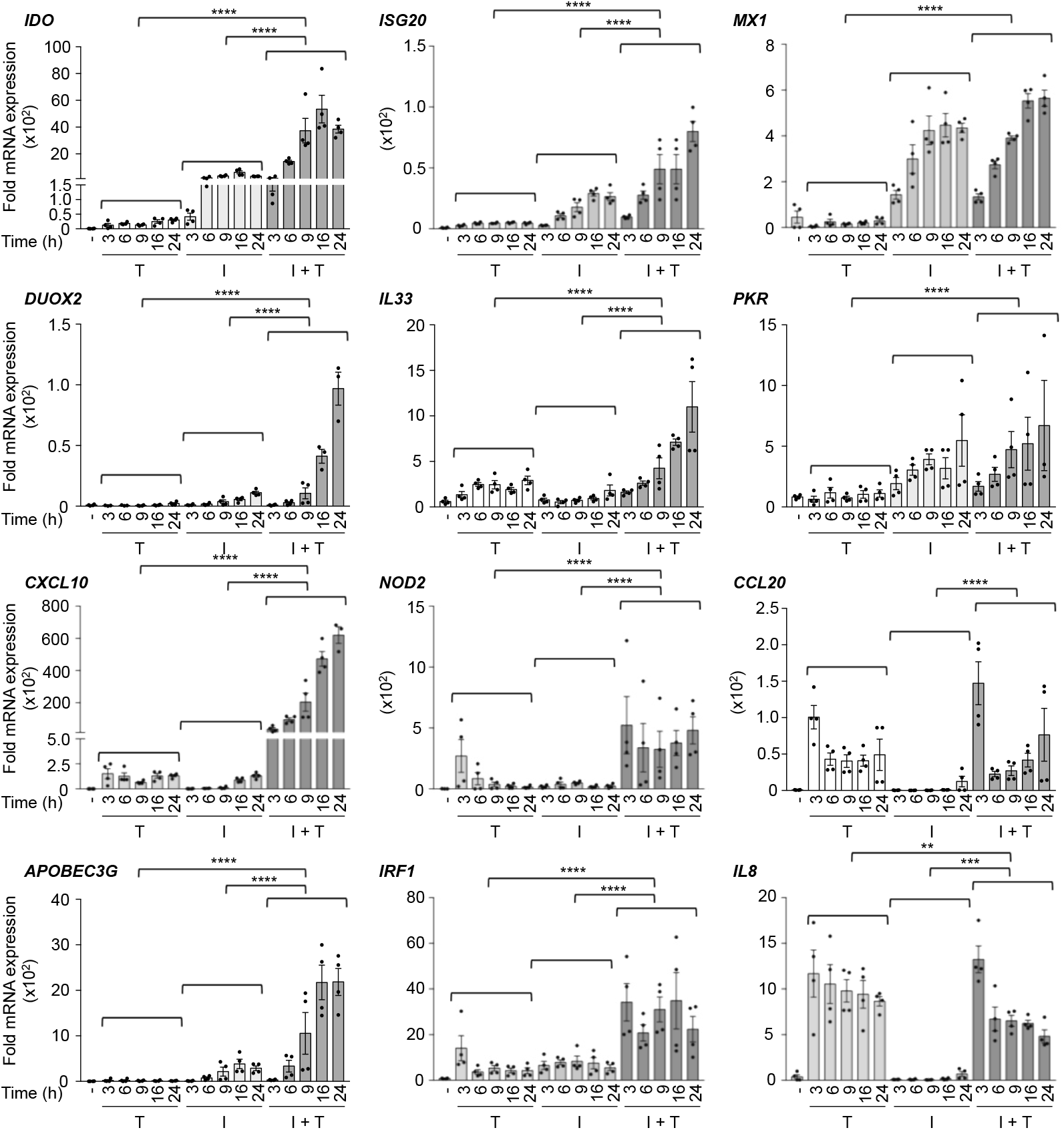
Expression of a panel of antiviral and immunoregulatory genes in response to IFNβ, TNFα and IFNβ+TNFα. A549 cells were stimulated with either TNFα, IFNβ or costimulated with IFNβ+TNFα for the indicated times. Quantification of mRNA was performed by qRT-PCR and expressed as fold expression after normalization to the S9 mRNA levels using the ΔΔCt method. Mean +/-SEM, n ≥3. Statistical comparison of TNFα *vs.* IFNβ+TNFα and IFNβ *vs.* IFNβ+TNFα was conducted using two-way ANOVA with Tukey’s post-test. *P* < 0.01 (**), *P* < 0.001 (***) or *P* < 0.0001 (****).

### Workflow for genome-wide characterization of the delayed transcriptional program induced by IFNβ+TNFα in the absence of STAT1

In a previous report, we provided evidence of a STAT1-independent, but STAT2- and IRF9-dependent, pathway engaged downstream of IFNβ+TNFα ^12^. Here, taking advantage of the STAT1-deficient human U3A cell line ^13^, we aimed to fully characterize the STAT1-independent transcriptional program induced by IFNβ+TNFα. Two hallmarks of STAT2 and IRF9 activation, *i.e.* STAT2 Tyr690 phosphorylation and induction of IRF9, were observed in U3A cells following stimulation with IFNβ+TNFα. Although STAT2 and IRF9 activation was reduced compared to the parental 2ftGH cells expressing endogenous STAT1, this observation supports the capacity of STAT2 and IRF9 to be activated in STAT1-deficient U3A cells stimulated with IFNβ+TNFα (**Figure 2A**).

**Figure 2:**
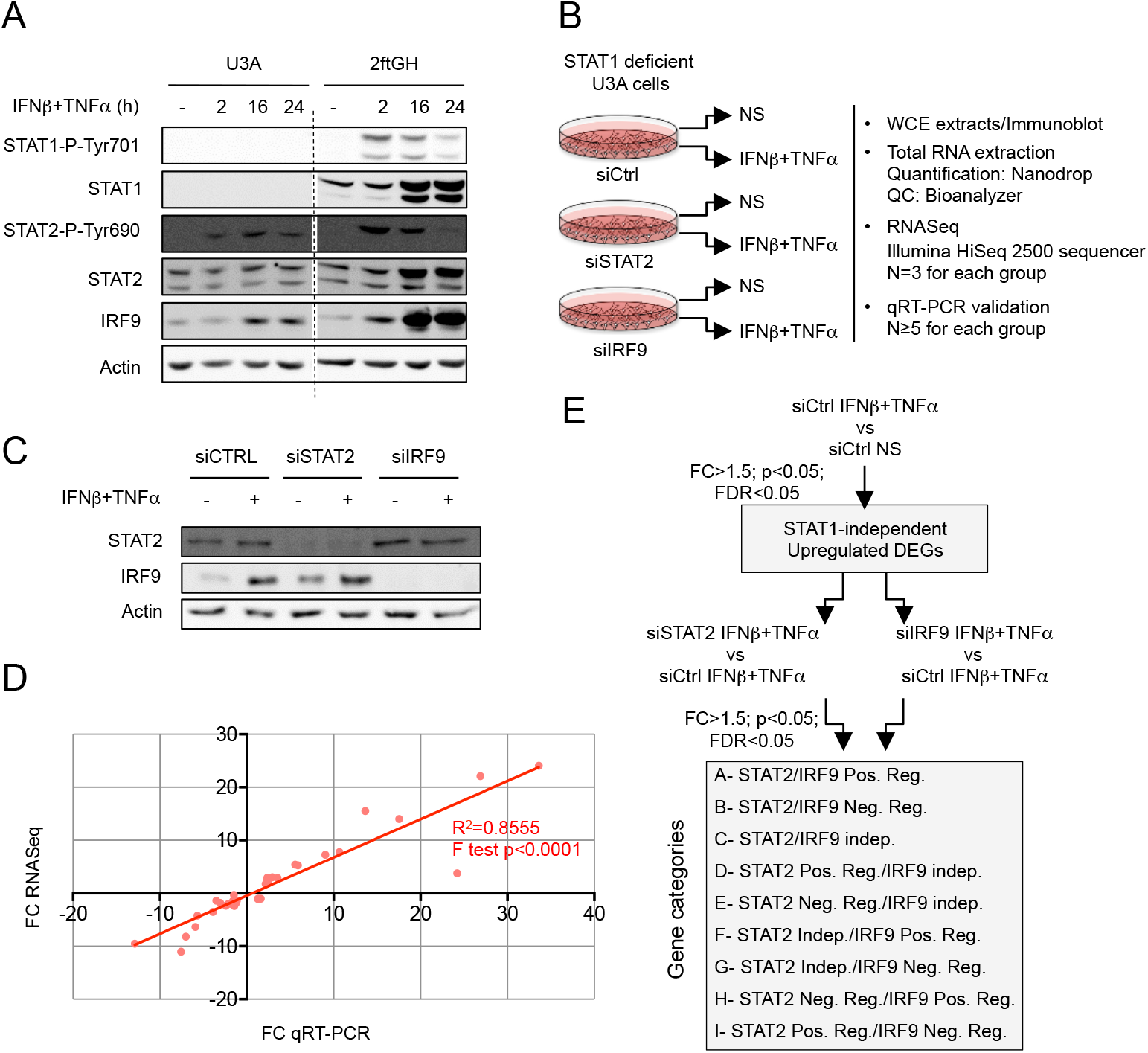
Experimental design used to study the STAT1-independent delayed transcriptional program induced by the combination of IFNβ and TNFα. **A)** U3A (STAT1-deficient) and 2ftGH (parental STAT1-positive) cells were left untreated or stimulated with IFNβ+TNFα for the indicated times. WCE were analyzed by SDS-PAGE followed by immunoblot using anti STAT1-P-Tyr701, total STAT1, STAT2-P-Tyr690, total STAT2, IRF9 or actin antibodies. **B-E)** U3A cells were transfected with siCTRL, siSTAT2 or siIRF9 before being left untreated (NS) or stimulated with IFNβ+TNFα for 24h. **In B**, The schematic describes the workflow of sample preparation and analysis. **In C**, WCE were analyzed by SDS-PAGE followed by immunoblot using anti STAT2, IRF9 and actin antibodies. **In D**, Graph showing the correlation between fold-changes (FC) measured by RNASeq and qRT-PCR for 13 genes. Data from siCTRL NS vs siCTRL IFNβ+TNFα, siSTAT2 IFNβ+TNFα vs siCTRL IFNβ+TNFα, siIRF9 IFNβ+TNFα vs siCTRL IFNβ+TNFα conditions were used. **In E**, Diagram describing the bioinformatics analysis strategy used to determine STAT1-independent differentially expressed genes (DEGs) and their regulation by STAT2 and IRF9.

To investigate the transcriptional program triggered independently of STAT1, the U3A cells were efficiently transfected with Control (Ctrl)-, STAT2- or IRF9-RNAi and further left untreated or stimulated with IFNβ+TNFα for 16h (**Figure 2B**). Efficient silencing was confirmed by immunoblot (**Figure 2C**). To perform genome-wide transcriptional analysis, total RNA was isolated and analyzed by RNA sequencing (n=3 for each group detailed in **Figure 2B**) on an Illumina HiSeq2500 platform. To validate the RNASeq data, the fold changes (FC) of 13 genes between the different experimental groups, siCTRL non-stimulated (NS) *vs* siCTRL IFNβ+TNFα, siCTRL IFNβ+TNFα *vs* siSTAT2 IFNβ+TNFα and siCTRL IFNβ+TNFα *vs* siIRF9 IFNβ+TNFα was confirmed by qRT-PCR. A positive linear relationship between RNASeq and qRT-PCR FC was observed (**Figure 2D**).

### An antiviral and immunoregulatory transcriptional signature is induced by IFNβ+TNFα independently of STAT1

To identify STAT1-independent differentially expressed genes (DEGs) upon IFNβ+TNFα stimulation, comparison between non-stimulated (NS) and IFNβ+TNFα stimulated control cells was performed (**Figure 2E**). In total, 612 transcripts, including protein-coding transcripts, pseudogenes and long non-coding RNA (lncRNA), were significantly different (FC>1.5, p<0.05, FDR<0.05) in IFNβ+TNFα *vs* NS. Among these, 482 DEGs were upregulated and 130 were downregulated (**Figure 3A;** See **Supplemental Table S1** for a complete list of DEGs). To identify the Biological Processes (BP) and Molecular Functions (MF) induced by IFNβ+TNFα independently of STAT1, we first subjected upregulated DEGs through Gene Ontology (GO) enrichment analysis. The top enriched GO BP (p< 10^-10^) and MF, are depicted in **Figure 3B (**See **Supplemental Table S2** for a complete list of enriched GO). The majority of the top enriched BPs were related to cytokine production and function (response to cytokine, cytokine-mediated signaling pathway, cytokine production, regulation of cytokine production), immunoregulation (Immune response, immune system process, innate immune response, regulation of immune system process) and host defense response (defense response, response to other organism, 2’-5’-oligoadenylate synthetase activity and dsRNA binding). Fourteen MF categories were found enriched in IFNβ+TNFα. The top enriched MFs were related to cytokine and chemokine functions (cytokine activity, cytokine receptor binding, chemokine activity, Interleukin 1-receptor binding). Other enriched MF included peptidase related functions (endopeptidase inhibitor activity, peptidase regulator activity, serine-type endopeptidase activity).

**Figure 3:**
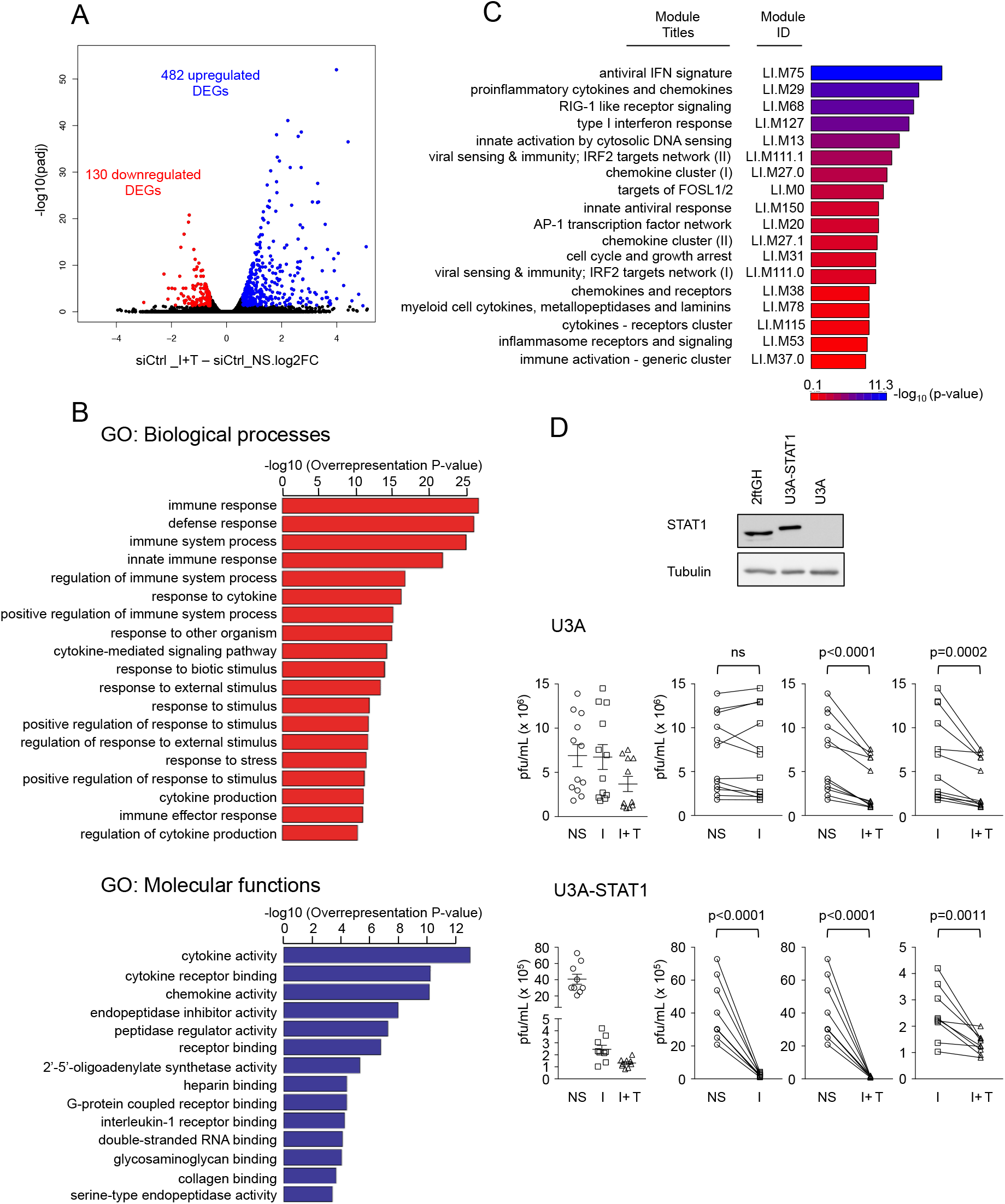
Functional characterization of STAT1-independent IFNβ+TNFα-induced DEGs. **A)** Volcano plots of the fold-change (FC) *vs.* adjusted *P*-value of IFNβ+TNFα (I+T) vs non-stimulated (NS) siCtrl conditions. **B)** Gene ontology (GO) enrichment analysis of the differentially upregulated genes in IFNβ+TNFα vs non-stimulated siCtrl conditions based on the Biological Processes and Molecular Function categories. **C)** Modular transcription analysis of upregulated DEGs. Eighteen enriched modules are shown. The full list of enriched modules is available **Supplemental Table S3. D)** U3A and STAT1-rescued U3A-STAT1 cells were stimulated with IFNβ (I) or IFNβ+TNFα for 30h before infection with VSV at a MOI of 5 for 12h. The release of infectious viral particles was quantified by plaque forming unit (pfu) assay. The left graphs show dot-plots of all stimulations. Statistical comparisons were performed on the “before and after” plots (displayed on the right of dot-plot graphs) using ratio paired t-tests. Top panel shows the immunoblot in 2ftGH, U3A-STAT1 and U3A cells using anti-STAT1 and anti-tubulin (loading control) antibodies.

To further gain insight into the functional significance of the STAT1-independent IFNβ+TNFα-induced transcriptional response, we conducted a modular transcription analysis of upregulated DEGs against 606 immune-related functional modules. These modules were previously defined from co-clustered gene sets built via an unbiased data-driven approach as detailed in the material and methods section ^14, 15^. STAT1-independent IFNβ+TNFα-induced DEGs showed significant enrichment in 37 modules (**Supplemental Table S3**). Six of these modules were associated with virus sensing/Interferon antiviral response, including LI.M75 (antiviral IFN signature), LI.M68 (RIG-I-like receptor signaling), LI.M127 (type I interferon response), LI.M111.0 and LI.M111.1 (IRF2 target network) and LI.M150 (innate antiviral response) (**Figure 3C**). Additionally, 6 modules were associated with immunoregulatory functions, including LI.M29 (proinflammatory cytokines and chemokines), LI.M27.0 and LI.M27.1 (chemokine cluster I and II), LI.M38 (chemokines and receptors), LI.M115 (cytokines receptors cluster) and LI.M37.0 (immune activation - generic cluster) (**Figure 3C**). Module analysis also showed enriched AP-1 transcription factor-related network modules, LI.M20 and LI.M0, and cell cycle and growth arrest LI.M31 module. To confirm the capacity of IFNβ+TNFα to trigger an antiviral response independently of STAT1, U3A cells were stimulated with IFNβ or IFNβ+TNFα and further infected with Vesicular stomatitis Virus (VSV). While VSV replicated similarly in untreated U3A cells and in cells treated with IFNβ, treatment with IFNβ+TNFα significantly restricted VSV replication (**Figure 3D**). Importantly, IFNβ treatment led to a significant antiviral response when STAT1 expression was restored in U3A cells (U3A-STAT1). In this context, the combination of IFNβ+TNFα was more effective in restricting VSV than IFNβ alone (**Figure 3D**). Collectively, these data demonstrate that the co-stimulation with IFNβ and TNFα induces a broad antiviral and immunoregulatory transcriptional program that is independent of STAT1.

### Clustering of STAT1-independent IFNβ+TNFα induced DEGs according to their regulation by STAT2 and IRF9

Next, we sought to assess how the STAT1-independent IFNβ+TNFα-induced antiviral and immunoregulatory transcriptional response was regulated by STAT2 and IRF9. To do so, we compared transcripts levels in siSTAT2_IFNβ+TNFα *vs* siCTRL_IFNβ+TNFα and siIRF9_IFNβ+TNFα *vs* siCTRL_IFNβ+TNFα conditions (**Figure 2E** and **Supplemental Table S1**). Volcano plots revealed that a fraction of IFNβ+TNFα-induced DEGs were significantly (FC>1.5, p<0.05, FDR<0.05) downregulated or upregulated upon silencing of STAT2 (**Figure 4A**) or IRF9 (**Figure 4B**). Nine distinct theoretical categories of DEGs can be defined based on their potential individual behavior across siSTAT2 and siIRF9 groups (Categories A-I, **Figure 2E**): a gene can either be downregulated upon STAT2 and IRF9 silencing, indicative of a positive regulation by STAT2 and IRF9 (Categorie A); conversely, a gene negatively regulated by STAT2 and IRF9 would exhibit upregulation upon STAT2 and IRF9 silencing (Categorie B); Genes that do not exhibit significant differential expression in siSTAT2 and siIRF9 groups would be classified as STAT2 and IRF9 independent (Categorie C); IRF9-independent genes could exhibit positive (Categorie D) or negative (Categorie E) regulation by STAT2; conversely, STAT2-independent genes might be positively (Categorie F) or negatively (Category G) regulated by IRF9; lastly, STAT2 and IRF9 could have opposite effects on a specific gene regulation (Categorie H and I). Based on *a priori* clustering of RNASeq data (**Figure 4C**) and analysis of the expression of 18 genes by qRT-PCR (**Figure 4D**), we found that STAT1-independent IFNβ+TNFα-induced DEGs clustered into only 7 of the 9 possible categories. Amongst the 482 DEGs, 163 genes exhibited either inhibition or upregulation following silencing of STAT2 and/or IRF9 (Category B-G). A large majority of upregulated DEGs, i.e. 319 out of the 482 DEGs, were not significantly affected by either STAT2 or IRF9 silencing, and were therefore classified as STAT2/IRF9-independent (**Figure 4C**). No genes were found in categorie H and only one gene was found in categorie I.

**Figure 4:**
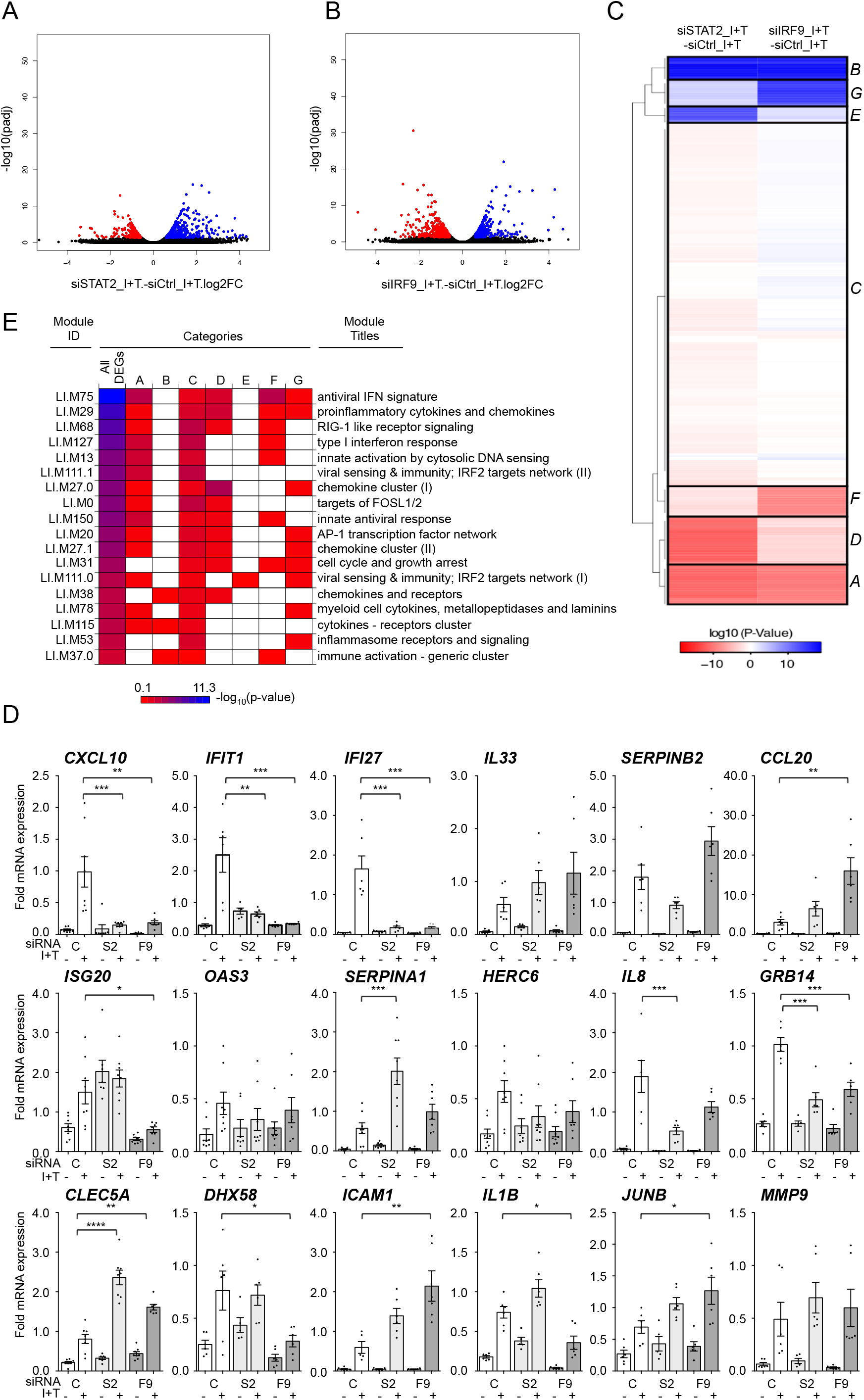
Clustering of IFNβ+TNFα-induced DEGs according to their regulation by STAT2 and IRF9. **A)** Volcano plots of the fold-change *vs.* adjusted *p*-value of siSTAT2 IFNβ+TNFα *vs.* siCTRL IFNβ+TNFα (I+T) conditions. **B)** Volcano plots of the fold-change *vs.* adjusted *p*-value of siIRF9 IFNβ+TNFα *vs.* siCTRL IFNβ+TNFα conditions. **C)** Hierarchical clustering of the categories of DEG responses according to their regulation by STAT2 and IRF9. Euclidean distance metric is used for the construction of distance matrix and the categories are used *a priori* input into clustering algorithm as detailed in Materials and Methods. **D)** Validation of DEGs regulation profile by qRT-PCR. U3A cells were transfected with siCTRL (C), siSTAT2 (S2) or siIRF9 (F9). Cells were further left untreated or stimulated with IFNβ+TNFα for 24h. Quantification of the mRNA corresponding to the indicated genes was performed by qRT-PCR and expressed as relative expression (ΔΔCt) after normalization to the S9 mRNA levels. Mean +/-SEM, n≥5. Statistical comparison of siSTAT2 IFNβ+TNFα *vs* siCTRL IFNβ+TNFα and siIRF9 IFNβ+TNFα *vs* siCTRL IFNβ+TNFα conditions was conducted using one-way ANOVA with Dunett post-test. *P* < 0.05 (*), *P* < 0.01 (**), *P* < 0.001 (***) or *P* < 0.0001 (****). **E)** Diagram showing enriched transcription modules in each gene category.

To functionally interpret these clusters, we applied the modular transcription analysis to each of the categories to assess for the specific enrichment of the functional modules found associated with IFNβ+TNFα-upregulated DEGs (**Figure 4E**). First, most modules, except LI.M31 (cell cycle and growth arrest), LI.M38 (chemokines and receptors), LI.M37.0 (immune activation - generic cluster) and LI.M53 (inflammasome receptors and signaling), were enriched in the category of DEGs positively regulated by STAT2 and IRF9 (Category A). Conversely, the cluster negatively regulated by STAT2 and IRF9 (Category B) exclusively contains enriched LI.M38 (chemokines and receptors), LI.M37.0 (immune activation - generic cluster) and LI.M115 (cytokines receptors cluster). The cluster of IRF9-independent genes that are negatively regulated by STAT2 (Category E) only exhibited enrichment in the virus sensing/ IRF2 target network LI.M111.0 module, while the IRF9-independent/ STAT2-positively regulated cluster (Category D) encompasses antiviral and immunoregulatory functions. The STAT2-independent, but IRF9 positively regulated transcripts (Category F) was mainly enriched in modules associated with the IFN antiviral response, including LI.M75 (antiviral IFN signature), LI.M68 (RIG-I-like receptor signaling), LI.M127 (type I interferon response), and LI.M150 (innate antiviral response). Finally, the STAT2-independent, but IRF9 negatively regulated cluster (Category G) was mostly enriched in modules associated with immunoregulatory functions, including LI.M29 (proinflammatory cytokines and chemokines), LI.M27.0 and LI.M27.1 (chemokine cluster I and II), LI.M78 (myeloid cell cytokines), but also with cell cycle and growth arrest (LI.M31) and inflammasome receptors and signaling (LI.M53). Of note, all modules were found enriched in the cluster of genes induced independently of STAT2 and IRF9 (Category C), pointing to a broad function of this yet to be defined pathway(s) in the regulation of the antiviral and immunoregulatory program elicited by IFNβ+TNFα. Altogether these observations reveal that STAT2 and IRF9 are involved in the regulation of only a subset of the genes induced in response to the co-stimulation by IFNβ and TNFα in the absence of STAT1. Importantly, our results also reveal that STAT2 and IRF9 act in a concerted manner to regulate a specific subset of the IFNβ+TNFα-induced DEGs, but are also independently involved in distinct non-canonical pathways.

### Differential regulation of *CXCL10* in response to IFNβ and IFNβ+TNFα

Identification of DEGs upregulated by IFNβ+TNFα in a STAT1-independent, but STAT2 and IRF9-dependent, manner potentially reflects the regulation by an alternative STAT2/IRF9-containing complex ^4, 7^. Whether this STAT2/IRF9 pathway ultimately leads to gene regulation through the same ISRE sites used by the ISGF3 complex remained to be assessed. In our RNASeq analysis (**Supplemental Table S1**) and qRT-PCR validation (**Figure 4D**), *CXCL10* belongs to the IFNβ+TNFα-induced DEGs that are dependent on STAT2 and IRF9. The *CXCL10* promoter contains 3 ISRE sites. We used full-length wild-type (972bp containing the 3 ISRE sites), truncated (376bp containing only the ISRE(3) site) or mutated (972bp containing a mutated ISRE(3) site) *CXCL10* promoter luciferase (*CXCL10*prom-Luc) reporter constructs (**Figure 5A**) to determine the ISRE consensus site(s) requirement. U3A and STAT1-rescued U3A-STAT1 cells were transfected with the *CXCL10*prom-Luc constructs and further stimulated with IFNβ or IFNβ+TNFα to monitor the canonical ISGF3-dependent and the STAT1-independent responses. In the absence of STAT1, IFNβ failed to activate *CXCL10*prom. However, the induction of the *CXCL10*prom activity was restored either upon expression of STAT1 or when TNFα was used together with IFNβ (**Figure 5B**). While the activation of *CXCL10*prom in response to IFNβ in the presence of STAT1 involves the distal ISRE(1) and/or ISRE(2) sites and the proximal ISRE(3) site, only the ISRE(3) site is required for induction by IFNβ+TNFα in the absence of STAT1. Additionally, we also assessed the contribution of the two NF-κB and the AP-1 sites present in the promoter using *CXCL10*prom-Luc mutated constructs in comparison to the wild-type reporter. While none of the NF-κB and the AP-1 sites were required for induction of the *CXCL10* promoter by IFNβ in the presence of STAT1, the two NF-κB sites were necessary for the STAT1-independent induction in response to IFNβ+TNFα (**Figure 5B**). These observations suggest that the ISRE site usage is more restricted in the absence of STAT1 in the context of the co-stimulation with IFNβ TNFα than in the context of an ISGF3-dependent regulation.

**Figure 5:**
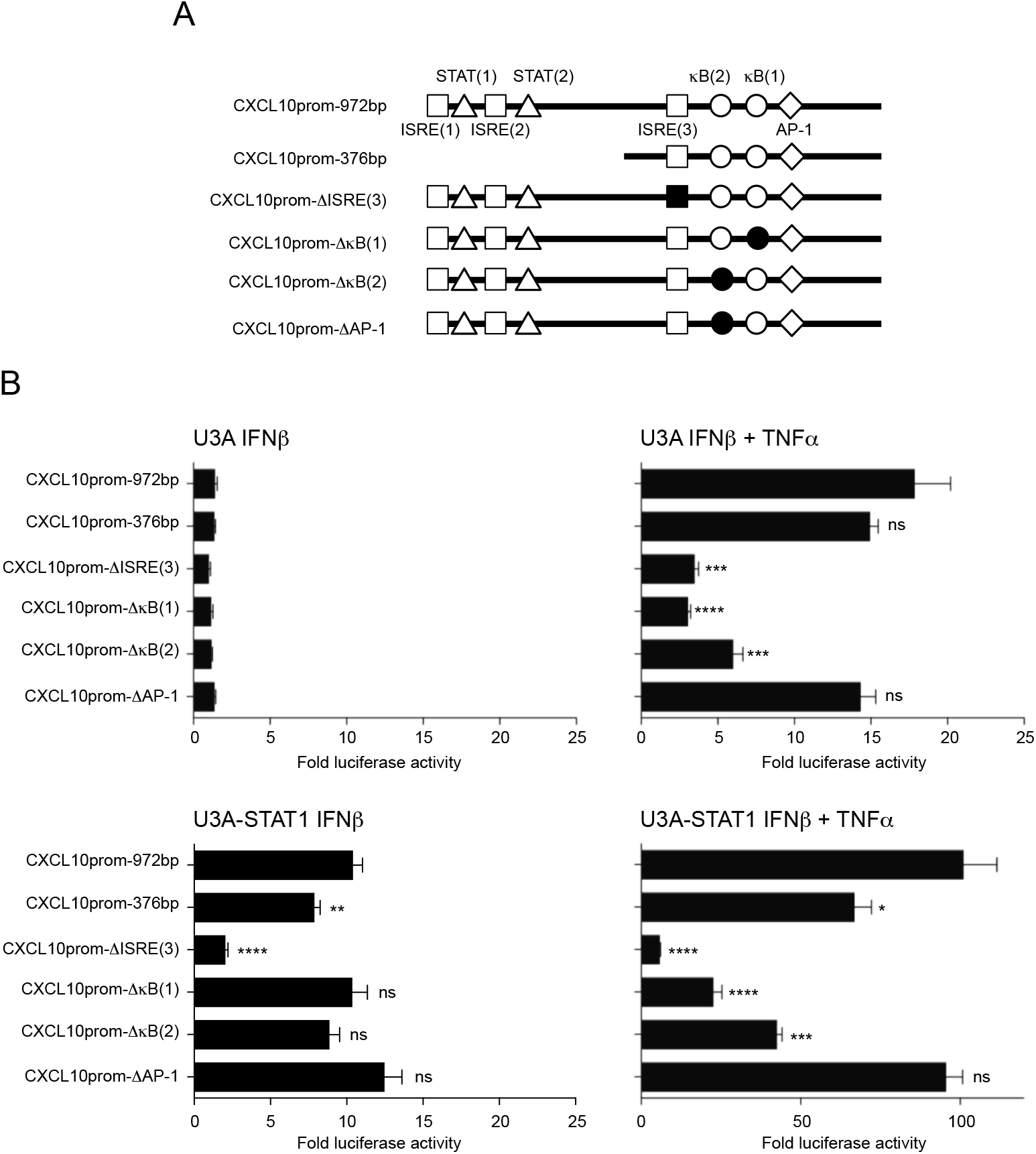
Analysis of the *CXCL10* promoter regulation in response to IFNβ+TNFα vs IFNβ. **A)** Schematic representation of the *CXCL10* promoter (CXCL10prom) luciferase constructs used in this study indicating the main transcription factors consensus binding sites. **B)** U3A and U3A-STAT1 cells were transfected with the indicated CXCL10prom-luciferase reporter constructs and either left untreated or stimulated with IFNβ or IFNβ+TNFα. Relative luciferase activities were measured at 16h post-stimulation and expressed as fold over the corresponding unstimulated condition. Mean +/-SEM, n=6. Statistical analyses were performed using an unpaired t-test comparing each promoter to the CXCL10prom-972bp construct. *P* < 0.01 (**), *P* < 0.001 (***).

## DISCUSSION

Our study was designed to determine the functional relevance of a STAT1-independent, but STAT2- and IRF9-dependent, signaling pathway in the transcriptional program induced by IFNβ in the presence of TNFα. Previous studies reported that IFNβ and TNFα synergize to induce a specific delayed antiviral program that differs from the response induced by IFNβ only. This specific synergy-dependent antiviral response is required for a complete abrogation of Myxoma virus in fibroblasts ^10^ and contributes to the establishment of a sustained type I and type III IFNs response during paramyxovirus infection in airway epithelial cells ^12^. The underlying mechanisms of this specific response remain ill defined, but their elucidation is of particular importance with regards to conditions with elevated levels of both IFNβ and TNFα such as pathogen intrusion or, autoimmune or chronic inflammatory diseases ^8^.

First, we confirmed the previously documented synergistic induction of an IFNβ+TNFα-mediated delayed transcriptional program composed of genes that are either not responsive to IFNβ or TNFα separately or are only responsive to either one of the cytokine when used separately albeit to a lesser extent (**Figure 1**). Using genome wide RNA sequencing, we demonstrate that IFNβ+TNFα induces a broad transcriptional program in cells deficient in STAT1. GO enrichment and transcriptional module analyses showed that STAT1-independent upregulated DEGs encompass a wide range of immunoregulatory and host defense, mainly antiviral, functions. The STAT1-independent antiviral response mounted by STAT1-deficient cells in response to IFNβ+TNFα efficiently restricted VSV replication (**Figure 2**). These findings highlight the functional significance of a STAT1-independent response.

We also focused on deciphering the role of STAT2 and IRF9 in the STAT1-independent transcriptional program elicited in response to IFNβ+TNFα. We previously found that IFNβ+TNFα induces the *DUOX2* gene via a STAT2- and IRF9-dependent pathway in the absence of STAT1 ^12^. To what extent this pathway contributes to the STAT1-independent response engaged downstream of IFNβ+TNFα remained to be addressed. Based on the possible regulation by STAT2 and/or IRF9, IFNβ+TNFα-induced DEGs could theoretically partitioned into 9 different predicted categories (**Figure 2E**), but we found that they in fact only significantly segregate into 7 categories. Importantly, the distribution of DEGs amongst 7 different categories reflects distinct contributions of STAT2 or IRF9 and highlights the heterogeneity of the mechanisms of regulation of the IFNβ+TNFα-induced genes. Importantly, only one anecdotic gene was found in categories implying inverse regulation by STAT2 and IRF9 (categories H and I) pointing to convergent functions of STAT2 and IRF9 when both are engaged in gene regulation. We can rule out that the distinct regulation mechanisms reflect distinct profiles of induction by IFNβ+TNFα. For instance *CXCL10*, *IL33*, *CCL20* and *ISG20* all exhibit synergistic induction by IFNβ+TNFα (**Figure 1**), but are differentially regulated by STAT2 and/or IRF9; while *CXCL10* is dependent on STAT2 and IRF9, *IL33* is independent on STAT2 and IRF9, and *CCL20* and *ISG20* are STAT2-independent but IRF9-dependent (**Figure 4D**).

Consistent with our previous report ^12^, we found several genes positively regulated by a non-canonical STAT2- and IRF9-dependent, but STAT1-independent, pathway (Category A). DEGs in this category encompass most of the functions induced in response to IFNβ+TNFα, with the notable exception of cell cycle and growth arrest and inflammasome and receptor signaling functions. Genes negatively regulated by STAT2 and IRF9 were also identified (Category B). Accumulating evidence point to the formation of an alternative STAT2/IRF9-containing complex mediating gene expression in the absence of STAT1 ^16-20^. The specificity of the DNA-binding of a STAT2/IRF9 complex compared to the ISGF3 complex remains unclear. It was originally found that STAT2/IRF9 exhibit only limited DNA-binding affinity for the typical ISRE sequence in the absence of STAT1 ^16^, but association of STAT2 with the promoter of antiviral genes induced by Dengue virus in STAT1-deficient mice was demonstrated by Chromatin immunoprecipitation ^21^. Here, the *CXCL10* gene was further analyzed as a paradigm of STAT2- and IRF9-positively regulated gene. Promoter activity analysis unveiled distinct ISRE site usage between IFNβ and IFNβ+TNFα stimulation (**Figure 5**). Interestingly, the ISRE site used in response to IFNβ+TNFα lies close to the NF-κB sites that are also specifically engaged in this response. A possible mechanism for the synergistic action of IFNβ+TNFα might be related to the previous description of the cooperativity between ISGF3 and NF-κB in the context of Listeria infection ^22, 23^. In this context, ISGF3 and NF-κB tether a complete functional mediator multi-subunit complex that bridges transcription factors with Pol II and initiation and elongation factors to the promoter of antimicrobial genes ^22^. Both STAT1 and STAT2 functionally and physically interact with the mediator ^24, 25^. Additionally, IL-6 induction by NF-κB inducers was found synergistically enhanced by IFNβ. In this model, unphosphorylated STAT2 was proposed to act as a bridge connecting p65 and IRF9 bound to κB and ISRE consensus sites, respectively ^26^. Therefore, in the absence of STAT1, the synergistic induction of some IFNβ+TNFα-induced DEGs might be explained by the interaction between a STAT2/IRF9-containing complex with NF-κB. However, we cannot exclude that interaction with a yet to be identified alternative transcription factor might tether STAT2 and IRF9 to the DNA in the absence of STAT1. Specific regulation of STAT2 and IRF9 might also contribute to the synergistic response elicited by IFNβ+TNFα. In a previous study, we observed IRF9 induction and enhanced/extended STAT2 phosphorylation in response to IFNβ+TNFα ^12^. We hypothesized that these events likely contribute to the specific activation of the non-canonical STAT2/IRF9-dependent pathway. Remarkably, a similar prolonged STAT2 phosphorylation was observed upon stimulation of STAT1 KO bone marrow–derived murine macrophages with IFNα. In this context, a STAT2/IRF9 complex was shown to induce a delayed set of ISGs ^20^. Taken together, it is reasonable to speculate that in a physiological context, when TNFα is present with IFNβ, a signal is elicited to progressively exclude STAT1 from the STAT2/IRF9 complex and favor the non-canonical STAT2/IRF9 pathway to drive a specific delayed response.

The observation that some IFNβ+TNFα-induced genes were independent of IRF9 but dependent on STAT2, either positively or negatively (Categories D and E), in a STAT1 deficient context might reflect the previous observation that STAT2 forms alternative complexes with other STAT members. STAT2 was shown to associate with STAT3 and STAT6, although it is not completely clear whether IRF9 is also required in these alternative complexes ^27, 28^. Interestingly, transcriptional module analyses demonstrated that the functional distribution of genes negatively regulated by STAT2 is very limited compared to other categories; only a virus-sensing module was enriched in this category. In contrary, IRF9-independent genes positively regulated by STAT2 mediate broader antiviral and immunoregulatory functions.

ISGF3-independent functions of IRF9 have been proposed based on the study of IRF9 deficiencies (reviewed in ^7, 29^). However, IRF9 target genes in these contexts have been barely documented. Intriguingly, Li et al. ^30^ studied IFNα-induced genes and their dependency on the ISGF3 subunits. While they confirmed previous studies showing that IFNα can trigger a delayed and sustained ISGs response via an ISGF3-independent pathway, it is very striking that they did not find STAT1- and STAT2-independent but IRF9-dependent genes. Indeed, all identified IRF9-dependent genes were either STAT2- or STAT1-dependent. This result greatly differs with our study. Here, we found several IFNβ+TNFα-induced DEGs independent of STAT1 and STAT2, but positively or negatively regulated by IRF9 (Categories F and G). Typically, IRF9 is considered a positive regulator of gene transcription. However, our findings are consistent with recent reports documenting the role of IRF9 in the negative regulation of the TRIF/NF-κB transcriptional response ^31^ or the expression of SIRT1 in acute myeloid leukemia cells ^32^. The molecular mechanism underlying gene regulation by IRF9 without association with either STAT1 or STAT2 remain to be elucidated. To the best of our knowledge, no alternative IRF9-containing complex has been described.

Unexpectedly, a vast majority of genes were found to be independent of STAT2 and IRF9 (Categorie C). All transcriptional modules were enriched in this category pointing to a major role of a yet to be identified pathway in the establishment of a host defense and immunoregulatory response. The NF-κB pathway is widely known to be engaged downstream of the TNF receptor. While NF-κB is an obvious candidate for being involved in the regulation of the STAT2 and IRF9-independent DEGs, this might fall short in explaining the synergistic action of IFNβ+TNFα. Synergistic activation of NF-κB was reported in the context of IFNγ and TNFα treatment ^33^. However, we did not observe enhanced NF-κB activation upon IFNβ+TNFα stimulation compared to TNFα alone ^12^. Alternative pathways might be of interest. For instance, the *IL33* gene is synergistically induced by IFNβ+TNFα in a STAT2 and IRF9-independent manner; scanning of the *IL33* promoter for transcription factor binding sites revealed AP-1, NF-κB and IRF7 consensus sites ^34^. The potential role of AP-1 is also supported by the finding that the AP-1 transcription network module is enriched amongst IFNβ+TNFα-induced DEGs (**Figure 4E**). However, this module is not restricted to genes regulated independently of STAT2 and IRF9. It is also worth noting that two modules of IRF2-target genes were enriched, although again not specifically in the STAT2- and IRF-9 independent category (**Figure 4E**). Finally, increased colocalized recruitment of IRF1 and p65 to the promoter of a subset of genes induced by IFNα and TNFα in macrophages was observed ^35^. However, while IRF1 was found synergistically induced by IFNβ+TNFα at early stages (**Figure 1**), we did not observe significant induction of IRF1 in the absence of STAT1 by RNASeq (**Supplemental Table S1**) or qRT-PCR (data not shown). Further studies will be needed to challenge the role of these various pathways in the synergistic induction of genes by IFNβ+TNFα independently of STAT2 and IRF9.

Altogether our results demonstrate that in conditions with elevated levels of IFNβ and TNFα, a broad antiviral and immunoregulatory delayed transcriptional program is elicited independently of STAT1. Our findings highlight the importance of diverse non-canonical STAT2 and/or IRF9 pathways. Consistent with the growing literature, IFNβ and TNFα synergistic action is in part mediated by the concerted action of STAT2 and IRF9, most likely present in an alternative complex. Finally, our study reveals specific independent roles of STAT2 and IRF9 in the regulation of distinct sets of IFNβ and TNFα-induced genes.

## MATERIAL AND METHODS

### Cell culture and stimulation

A549 cells (American Type Culture Collection, ATCC) were grown in F-12 nutrient mixture (Ham) medium supplemented with 10% heat-inactivated fetal bovine serum (HI-FBS) and 1% L-glutamine. The 2ftGH fibrosarcoma cell line and the derived STAT1-deficient U3A cell line, a generous gift from Dr. G. Stark, Cleveland, USA ^13^, were grown in DMEM medium supplemented with 10% HI-FBS or HI-Fetal Clone III (HI-FCl) and 1% L-glutamine. U3A cells stably expressing STAT1 were generated by transfection of the STAT1 alpha flag pRc/CMV plasmid (Addgene plasmid #8691; a generous gift from Dr. J. Darnell, Rockfeller University, USA ^36, 37^) and selection with 800μg/ml Geneticin (G418). Monoclonal populations of U3A stably expressing STAT1 cells were isolated. A pool of two clones, referred to as U3A-STAT1, was used in the experiments to mitigate the clonal effects. U3A-STAT1 cells were maintained in culture in DMEM supplemented with 10% HI-FCl, 1% Glu, and 200μg/ml G418. All cell lines were cultured without antibiotics. All media and supplements were from Gibco, with the expection of HI-FCl, which was from HyClone. Mycoplasma contamination was excluded by regular analysis using the MycoAlert Mycoplasma Detection Kit (Lonza). Cells were stimulated with IFNα (1000 U/mL, PBL Assay Science), TNFβ (10 ng/mL, R&D Systems) or IFNβ (1000 U/mL) +TNFα (10 ng/mL) for the indicated times.

### siRNA Transfection

The sequences of non-targeting control (Ctrl) and STAT2- and IRF9-directed RNAi oligonucleotides (Dharmacon, USA) have previously been described in ^12^. U3A cells at 30% confluency were transfected using the Oligofectamine transfection reagent (Life technologies). RNAi transfection was pursued for 48 h before stimulation.

### Immunoblot analysis

Cells were lysed on ice using Nonidet P-40 lysis buffer as fully detailed in ^38^. Whole-cell extracts (WCE) were quantified using the Bradford protein assay (Bio-Rad), resolved by SDS-PAGE and transferred to nitrocellulose membrane before analysis by immunoblot. Membranes were incubated with the following primary antibodies, anti-actin Cat #MAB1501 from Millipore, anti-IRF9 Cat #610285 from BD Transduction Laboratories, and anti-STAT1-P-Tyr701 Cat #9171, anti-STAT2-P-Tyr690 Cat #4441, anti-STAT1 Cat #9172, anti-STAT2 Cat #4594, all from Cell Signaling, before further incubation with horseradish peroxidase (HRP)-conjugated secondary antibodies (KPL or Jackson Immunoresearch Laboratories). Antibodies were diluted in PBS containing 0.5% Tween and either 5% nonfat dry milk or BSA. Immunoreactive bands were visualized by enhanced chemiluminescence (Western Lightning Chemiluminescence Reagent Plus, Perkin-Elmer Life Sciences) using a LAS4000mini CCD camera apparatus (GE healthcare).

### RNA isolation and qRT-PCR analyses

Total RNA was prepared using the RNAqueous-96 Isolation Kit (Ambion) following the manufacturer’s instructions. Total RNA (1μg) was subjected to reverse transcription using the QuantiTect Reverse Transcription Kit (Qiagen). Quantitative PCR were performed using either Fast start SYBR Green Kit (Roche) for *Mx1*, *IDO*, *APOBEC3G*, *CXCL10*, *NOD2*, *PKR*, *IRF1,* and *IFIT1* and *IL8* or TaqMan Gene Expression Assays (Applied Biosystems) for *DUOX2*, *IFI27*, *SERPINB2*, *IL33*, *CCL20*, *ISG20*, *OAS3*, *GRB14, CLEC5A, DGX58, ICAM1, IL1B, JUNB, MMP9, SERPINA1 and HERC6*. Sequences of oligonucleotides and probes used in PCR reactions are described in **Supplemental Table S4**. Data collection was performed on a Rotor-Gene 3000 Real Time Thermal Cycler (Corbett Research). Gene inductions were normalized over S9 levels, measured using Fast start SYBR Green Kit or TaqMan probe as necessary. Fold induction of genes was determined using the ΔΔCt method ^39^. All qRT-PCR data are presented as the mean ± SEM.

### RNA-sequencing (RNASeq)

Total RNA prepared as described above was quantified using a NanoDrop Spectrophotometer ND-1000 (NanoDrop Technologies, Inc.) and its integrity was assessed using a 2100 Bioanalyzer (Agilent Technologies). Libraries were generated from 250 ng of total RNA using the NEBNext poly(A) magnetic isolation module and the KAPA stranded RNA-Seq library preparation kit (Kapa Biosystems), as per the manufacturer’s recommendations. TruSeq adapters and PCR primers were purchased from IDT. Libraries were quantified using the Quant-iT™ PicoGreen^®^ dsDNA Assay Kit (Life Technologies) and the Kapa Illumina GA with Revised Primers-SYBR Fast Universal kit (Kapa Biosystems). Average size fragment was determined using a LabChip GX (PerkinElmer) instrument. Massively parallel sequencing was carried out on an Illumina HiSeq 2500 sequencer. Read counts were obtained using HTSeq. Reads were trimmed from the 3’ end to have a Phred score of at least 30. Illumina sequencing adapters were removed from the reads and all reads were required to have a length of at least 32bp. Trimming and clipping was performed using Trimmomatic ^40^. The filtered reads were aligned to the Homo-sapiens assembly GRCh37 reference genome. Each readset was aligned using STAR ^41^ and merged using Picard (http://broadinstitute.github.io/picard/). For all samples, the sequencing resulted in more than 29 million clean reads (ranging from 29 to 44 million reads) after removing low quality reads and adaptors. The reads were mapped to the total of 63679 gene biotypes including 22810 protein-coding genes. The non-specific filter for 1 count-per million reads (CPM) in at least three samples was applied to the reads and 14,254 genes passed this criterion. The entire set of RNAseq data has been submitted to the Gene Expression Omnibus (GEO) database (http://www.ncbi.nlm.nih.gov/geo) under accession number GSE111195.

### Bioinformatics analysis

Differential transcripts analysis. A reference-based transcript assembly was performed, which allows the detection of known and novel transcripts isoforms, using Cufflinks ^42^, merged using Cuffmerge (cufflinks/AllSamples/merged.gtf) and used as a reference to estimate transcript abundance and perform differential analysis using Cuffdiff and Cuffnorm tool to generate a normalized data set that includes all the samples. FPKM values calculated by Cufflinks were used as input. The transcript quantification engine of Cufflinks, Cuffdiff, was used to calculate transcript expression levels in more than one condition and test them for significant differences. To identify a transcript as being differentially expressed, Cuffdiff tests the observed log-fold-change in its expression against the null hypothesis of no change (i.e. the true log-fold-change is zero). Because of measurement errors, technical variability, and cross-replicate biological variability might result in an observed log-fold-change that is non-zero, Cuffdiff assesses significance using a model of variability in the log-fold-change under the null hypothesis. This model is described in details in ^43^. The differential gene expression analysis was performed using DESeq ^44^ and edgeR ^45^ within the R Bioconductor packages. Genes were considered differentially expressed between two group if they met the following requirement: fold change (FC) > ±1.5, p<0.05, FDR <0.05.

Enrichment of gene ontology (GO). GO enrichment analysis amongst differentially expressed genes (DEGs) was performed using Goseq ^46^ against the background of full human genome (hg19). GO-terms with adjusted p value < 0.05 were considered significantly enriched.

Clustering of DEGs. We categorized the DEGs according to their response upon silencing of siSTAT2 and siIRF9; categories are listed as A to I (Figure 2E). Then to determine relationship between these categories, we calculated the distance of centers of different categories. For each gene, we transformed siSTAT2 and siIRF9 fold changes (FC) to deviation from the mean FC of the category the respective gene belongs: *FC_new_ = FC_old_ - ε (FC_category_)*. The parameter *ε* was estimated to give the perfect match between predefined categories (A to I) and clustering based on Euclidean distance. Results were plotted as a heatmap.

Modular transcription analysis. The *tmod* package in R ^47^ was used for modular transcription analysis. In brief, each transcriptional module is a set of genes, which shows coherent expression across many biological samples ^48, 49^. Modular transcription analysis then calculates significant enrichment of a set of foreground genes, here DEGs, in pre-defined transcriptional module compared to a reference set. For transcriptional modules, we used a combined list of 606 distinct functional modules encompassing 12712 genes, defined by Chaussabel et al. ^14^ and Li et al. ^15^, as the reference set in tmod package (**Supplemental Table S5**). The hypergeometric test devised in *tmodHGtest* was used to calculate enrichments and p-values employing Benjamini-Hochberg correction ^50^ for multiple sampling. All the statistical analyses and graphical presentations were in performed in R ^51^.

### Luciferase gene reporter assay

U3A or U3A-STAT1 cells at 90% confluency were cotransfected with 100 ng of one of the following CXCL10 promoter containing firefly luciferase reporter plasmids (generously donated by Dr. David Proud, Calgary, ^52^), CXCL10prom-972pb-pGL4 (full length −875/+97 promoter), CXCL10prom-376pb-pGL4 (truncated −279/+97 promoter), CXCL10prom972pb-ΔISRE(3)-pGL4 (full length promoter with ISRE(3) site mutated), CXCL10prom972pb-ΔκB(1)-pGL4 (full length promoter with NFκB(1) site mutated), CXCL10prom972pb-ΔκB(2)-pGL4 (full length promoter with NFκB(2) site mutated), CXCL10prom972pb-ΔAP-1-pGL4 (full length promoter with AP-1 site mutated) together with 50ng of pRL-null renilla-luciferase expressing plasmid (internal control). Transfection was performed using Lipofectamine 2000 (Life technologies) using a 1:2 DNA to lipofectamine ratio. At 8 h post-transfection, cells were stimulated for 16 h with either IFNβ or IFNβ+TNFα. Firefly and renilla luciferase activities were quantified using the Dual-luciferase reporter assay system (Promega). Luciferase activities were calculated as the luciferase/renilla ratio and were expressed as fold over the non-stimulated condition.

### Virus titration by plaque assay

Quantification of VSV infectious virions was achieved through methylcellulose plaque forming unit assays. U3A and U3A-STAT1 cells were either left untreated or stimulated with IFNβ or IFNβ+TNFα for 30h. Cells were then infected with Vesicular Stomatitis Virus (VSV)-GFP (kindly provided by Dr. J. Bell, University of Ottawa, Canada) at an MOI of 5 for 1h in serum free medium (SFM). Cells were then washed twice with SFM and further cultured in DMEM medium containing 2% HI-FCl. The supernatants were harvested at 12h post-infection and serial dilutions were used to infect confluent Vero cells (ATCC) for 1h in SFM. The medium was then replaced with 1% methylcellulose in DMEM containing 10% HI-FCl. Two days post-infection, GFP-positive plaques were detected using a Typhoon Trio apparatus and quantified using the Imagequant software (Molecular Dynamics).

### Statistical analyses

Statistical analyses of qRT-PCR and luciferase assay results were performed using the Prism 7 software (GraphPad) using the tests indicated in the figure legends. Statistical significance was evaluated using the following *P* values: *P* < 0.05 (*), *P* < 0.01 (**), *P* < 0.001 (***) or *P* < 0.0001 (****). Differences with a *P*-value < 0.05 were considered significant. Statistical analysis of the RNASeq data is described in the Bioinformatics analysis section above.

## ACKNOWLEDGEMENTS

We are very thankful to Dr. D. Proud (University of Calgary, Canada), Dr. G. Stark (Cleveland clinic, USA) and Dr. J. Bell (University of Ottawa, Canada) for sharing reagents used in this study. RNASeq analyses were performed at the McGill University and Génome Québec Innovation Centre. The present work was funded by grants from Natural Sciences and Engineering Research Council of Canada (NSERC RGPIN/355306-2012 and RGPIN/2018-04279) and by the Research Chair in signaling in virus infection and oncogenesis from the Université de Montréal to NG. AS acknowledges funds from the University of Montreal, NSERC and the Merck Foundation. SCO was the recipient of a MITACS Globalink studentship. NG was recipient of a Tier II Canada Research Chair.

## AUTHOR CONTRIBUTIONS

MM and NG conceived and designed the experiments. MKM, AF, EC, MK, ANH, DK, SCO, and EM performed experiments. PD and AS performed bioinformatics analysis. MKM and NG analyzed the data. NG wrote the manuscript. All co-authors edited and approved the manuscript.

